# Single-Cell Profiling Reveals a Naive-Memory Relationship between CD56^bright^ and Adaptive Human Natural Killer Cells

**DOI:** 10.1101/2023.09.23.559062

**Authors:** M. Kazim Panjwani, Simon Grassmann, Rosa Sottile, Jean-Benoît Le Luduec, Theodota Kontopoulos, Kattria van der Ploeg, Joseph C. Sun, Katharine C. Hsu

## Abstract

Development of antigen-specific memory upon pathogen exposure is a hallmark of the adaptive immune system. While natural killer (NK) cells are considered part of the innate immune system, humans exposed to the chronic viral pathogen cytomegalovirus (CMV) often possess a distinct NK cell population lacking in individuals who have not been exposed, termed “adaptive” NK cells. To identify the “naïve” population from which this “memory” population derives, we performed phenotypic, transcriptional, and functional profiling of NK cell subsets. We identified immature precursors to the Adaptive NK cells that are equally present in both CMV+ and CMV-individuals, resolved an Adaptive transcriptional state distinct from most mature NK cells and sharing a common gene program with the immature CD56^bright^ population, and demonstrated retention of proliferative capacity and acquisition of superior IFNγ production in the Adaptive population. Furthermore, we distinguish the CD56^bright^ and Adaptive NK populations by expression of the transcription factor CXXC5, positioning these memory NK cells at the inflection point between innate and adaptive lymphocytes.

## Introduction

The adaptive immune system has been classically defined by two features: expression of a unique antigen receptor, and the ability to form immunological memory^1^. Conferring the ability of an immune cell to respond to a specific pathogen more quickly and robustly upon subsequent challenges following primary exposure, memory is linked to the antigen receptor for teleological and technical reasons: endowing antigen specificity to the recall response and enabling the tracing of a single naïve clone’s progeny following antigen exposure. However, the hallmark features of immunological memory - rapid and robust effector function, retention of proliferative capacity, and longevity - are grounded in the basic biology of a lymphocyte, and not just the expression of a rearranged protein sequence.

The contrast between the biology and identification of immune memory is best demonstrated by the classification of immune cells: all adaptive immune cells are lymphocytes, but not all lymphocytes are adaptive immune cells, traditionally defined by surface expression of an antigen-specific receptor. Lymphocytes not encompassed by the adaptive immune system include natural killer (NK) cells, which do not express a unique, rearranged antigen receptor on their cell surface and are therefore traditionally considered innate lymphocytes. The presence of an NK cell subset with superior function following certain viral exposures, however, contradicts this classification. In mice infected with mouse cytomegalovirus (MCMV), a population of NK cells can proliferate rapidly, clear virally infected cells, contract, and then perform these same functions again when expose to MCMV a second time^2^. In a subset of humans with prior exposure to human CMV, there is an outgrowth of a stable NK subpopulation with homogenous activating and inhibitory receptor expression that responds robustly to CMV-infected fibroblasts in vitro^3–5^. The distinction of NK cells from the adaptive immune cells on the basis of absence or presence of an antigen receptor is blurred by the expression of viral antigen-recognizing receptors found on some NK populations: in mice, Ly49H, which binds MCMV-derived m157 protein^6,7^; and in humans, NKG2C, which binds certain HCMV UL40-derived peptides presented on HLA-E^8,9^. In concession to their apparent capacity for expansion upon re-challenge with their respective cognate antigen, these receptor-expressing subsets are often referred to as “memory-like” or “adaptive” NK cells.

The identification of an Adaptive NK precursor, an analog to the naïve T lymphocyte, has been limited by the absence of a universally accepted phenotypic definition for the Adaptive NK population itself in humans. Various combinations of the expression of NKG2C, CD57, self– MHC-specific KIR, high levels of CD2, DNAM-1, or CD16, and the absence of FcRγ, NKp30, NKp46, PLZF, or EAT2 have all been used to define this population^3,5,10–13^. None of these features, however, is both exclusive to and necessary for Adaptive NK cells: NKG2C+ NK cells are present in CMV-seronegative (CMV-) individuals, and Adaptive NK cells have been found even in CMV+ individuals lacking the *KLRC2* gene encoding NKG2C^11^. Thus, attempts at identifying Adaptive NK cells through single-cell RNAseq have been stymied by the reliance upon *KLRC2* expression to identify the cells. Heterogeneity confounds the evaluation of the functional capacity of Adaptive and other NK cell subsets; NK cells can be stimulated through multiple receptors which can be preferentially expressed by different subsets, thus biasing the response to any single stimulus^14^. Furthermore, tracking of NK cells in response to infection in vivo is not routine in humans, hindering the identification of a precursor prior to exposure and tracing its differentiation into Adaptive and potentially other NK subsets.

Using a multimodal approach incorporating single-cell profiling at the transcript, protein, and functional level, we provide evidence that the relationship between a subset of the CD56^bright^ and Adaptive NK populations fits the naive-memory lymphocyte framework. We describe an immature precursor population present in pathogen-naive individuals, identify unique but tightly linked CD56^bright^ and Adaptive NK transcriptional states that are distinct from the classical CD56^dim^ NK population, and demonstrate the acquisition of priming-independent superior effector function and retention of proliferative capacity by the Adaptive NK population.

## Results

### Co-Expression of NKG2A and NKG2C on Immature NK Cells Independent of CMV Exposure

Whereas NKG2C is neither necessary nor sufficient to define Adaptive NK cells, the receptor does recognize a CMV-derived peptide and the frequency of NK cells expressing it increases after CMV exposure in individuals who possess the gene. Therefore, we used NKG2C as a beacon for Adaptive potential or identity, rather than marking the strict boundary of the population. Among human peripheral blood NK cells, we found that a minority of NKG2C+ NK cells are both negative for CD57, a marker for presumed terminal differentiation^15–17^, and positive for CD62L, a lymph organ homing molecule present on memory and naive T cells^18^ (Figure 1A). Intriguingly, the majority of these presumably immature NKG2C+ NK cells co-expressed the inhibitory receptor NKG2A, while the majority of CD57+ and/or CD62L-NKG2C+ NK cells did not (Figure 1B). While the presence of Adaptive NK cells is almost exclusive to individuals who are CMV+, a marker of CMV “antigen exposure”^3^, these NKG2C+NKG2A+ (A+C+) NK cells were found in equal frequency among both CMV+ and CMV-individuals when surveying 142 healthy donors (Figure 1C). The copy number of the *KLRC2* gene encoding NKG2C is correlated with A+C+ frequency, indicating that their population frequency is intrinsic to the host genotype and independent of antigen exposure (Figure S1A). The frequency of A+C+ NK cells is positively correlated with the frequency of A-C+ NK cells among CMV-individuals; this association is lost in CMV+ individuals in whom the A-C+ population has undergone significant expansion (Figure 1C). We have also found that A+C+ NK cell reconstitution precedes or is concurrent to A-C+ NK cell reconstitution in hematopoietic stem cell transplant recipients, making them a likely developmental prerequisite for the A-C+ NK cell population (Figure 1D)^19^.

**Figure 1.**
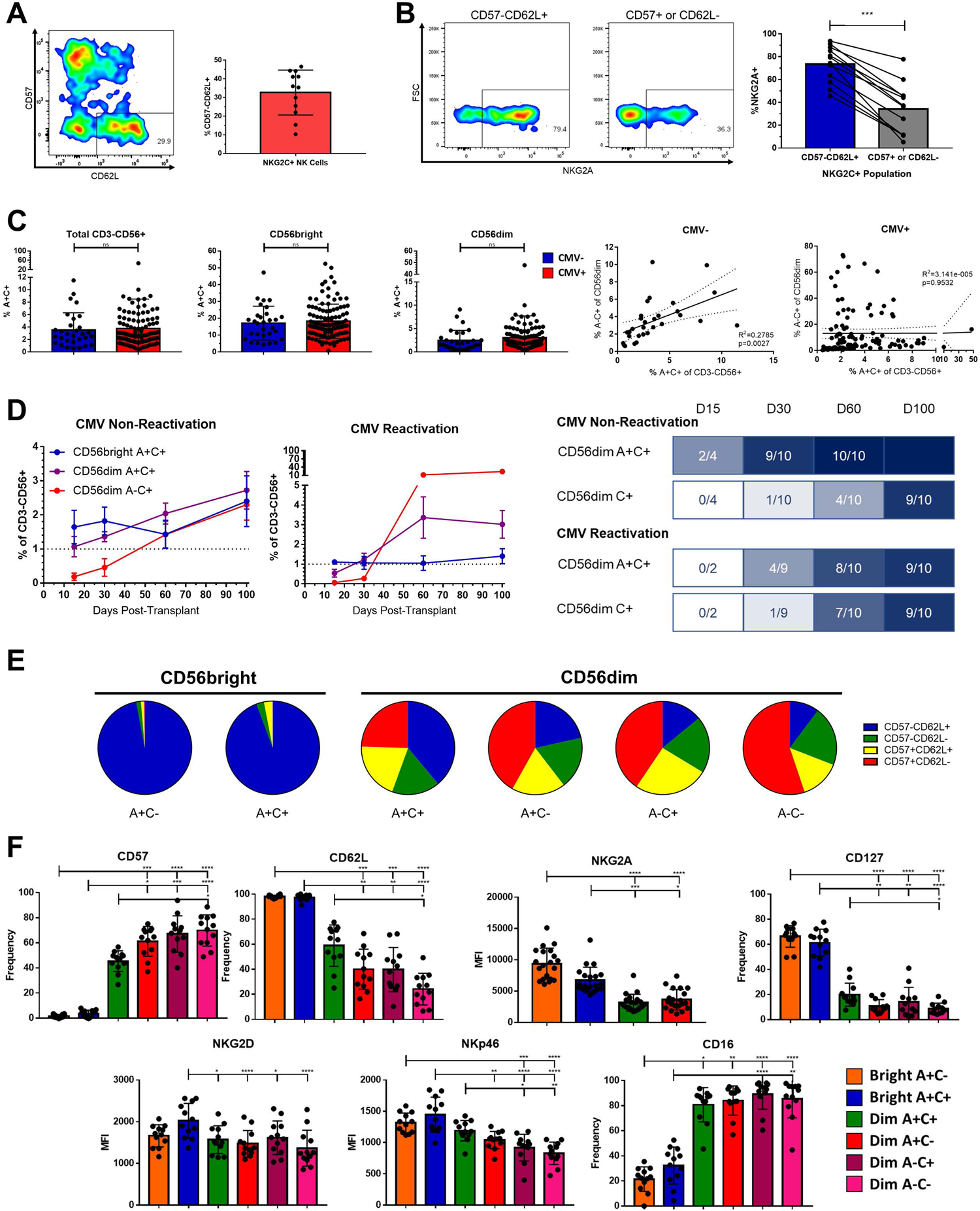
Phenotypic identification and characterization of an immature NKG2C+ precursor for Adaptive NK cells. A) Left, expression of CD57 and CD62L on live CD3-CD56+NKG2C+ NK cells in a representative healthy donor. Right, frequency of CD57-CD62L+ cells among NKG2C+ NK; bar represents mean and SD. B) Left, expression of NKG2A in NKG2C+ NK cell populations from a representative healthy donor. Right, frequency of NKG2A expression in NKG2C+ NK cell populations; lines connect paired donors, Wilcoxon matched pairs signed-rank test performed. C) Top, frequency of A+C+ among indicated populations from either CMV-seronegative or seropositive healthy donors; Mann-Whitney Test performed. Bottom, frequency of A+C+ and A-C+ among CD56^dim^ NK cells from CMV-seronegative or seropositive healthy donors; Spearman correlation coefficient shown. D) Left, mean frequency of populations among total NK cells in peripheral blood of HSCT recipients with and without CMV re-activation in this timespan. Right, timelines showing fraction of patients reaching reconstitution threshold of 1%. E) Proportions of CD57 and CD62L expression among NK cell subsets from 12 healthy donors; mean represented. F) Expression of cell surface markers on NK cell populations from healthy donors Dunn’s multiple comparison test performed, comparing groups below tick marks to group below capped end.

Although transient upregulation of NKG2A on mature NKG2C+ NK cells in the presence of IL-12 has been reported^20^, the phenotype of these A+C+ cells at baseline from healthy donors indicates they are not solely mature. A+C+ NK cells make up only 3.7% of the total CD3-CD56+ population but compose a substantial frequency of the CD56^bright^ population (18.0% vs 2.9% of the CD56^dim^). Like the CD56^bright^ A+C-cells, the CD56^bright^ A+C+ NK cells are almost entirely CD57-CD62L+, supporting their immature status (Figure 1E). Even among the CD56^dim^ NK cells, the A+C+ population is composed of a higher proportion of CD57-CD62L+ cells than the other 3 subsets of CD56^dim^ NK cells (A+C-, A-C+, A-C-). In addition to CD57 and CD62L, the A+C+ populations fill in a gradient between the CD56^bright^ and CD56^dim^ NK cell populations using other markers such as NKG2A, IL-7Rα (CD127), and CD16 (Figure 1F). The function of these A+C+ NK cells against tumor targets correlates with the expression of the cognate activating receptors: increased degranulation and IFNγ production against the NK-sensitive K562 and 721.221 lines, and intermediate response to ADCC of BE(2)n cells labeled with anti-GD2 antibody (Figure S1B). As the activating receptor NKG2C and inhibitory receptor NKG2A both recognize the HLA-E:peptide complex^21^, we challenged NK cells with K562 cells expressing equal amounts of HLA-E loaded with different peptides (Figure S1C). We found that inhibition from NKG2A was dominant, but the degree of inhibition appeared to be peptide-sensitive; among CD56^dim^ A+C+ and A+C-NK cells, in which the expression of NKG2A is similar, inhibition to HLA-E:G*01 was modestly mitigated in the NK cells co-expressing NKG2C, suggesting contribution from the activating receptor partially offsets the inhibition via NKG2A.

### Adaptive NK Cells Possess a Distinct Transcriptional Profile

As the A+C+ NK population appears to be a CMV-independent precursor to the Adaptive NK population, we sought to elucidate the relationship between these populations and the other peripheral blood NK cell population through single-cell RNAseq. NK cells from two CMV+ healthy donors were sorted according to expression of CD56, NKG2A, and NKG2C, labeled with corresponding hashtag oligos, and then recombined in equal numbers to overcome the rarity of some populations (Figure S2A). The CD56^dim^ A-C+ NK cells were split into a subset with two additional features of adaptiveness (NKp30-FcRγ-, Adaptive), and a subset with the inverse phenotype (NKp30hiFcRγ+, “non-Adaptive”) which is found also in CMV-individuals^3,22^. Data from the two donors were analyzed separately to avoid sex differences or weighting from a dominant donor, and consistent results between the donors were examined further.

The overall structure of the UMAP constructed from the transcriptomes is consistent between the two donors. Using the hashtag oligos to trace back the phenotypic origin of the individual cells, there are three major groupings: the CD56^bright^ A+C- and CD56^bright^ A+C+ group together; almost the entire A-C+ NKp30-Adaptive population is joined by half of the A-C+ NKp30hi “non-Adaptive” population to form the second group; and the other half of the A-C+ NKp30hi “non-Adaptive” A-C+ NK cells co-localizes with the remaining CD56^dim^ populations (A+C-, A+C+, and A-C-) for the third group (Figure 2A, S2B).

**Figure 2.**
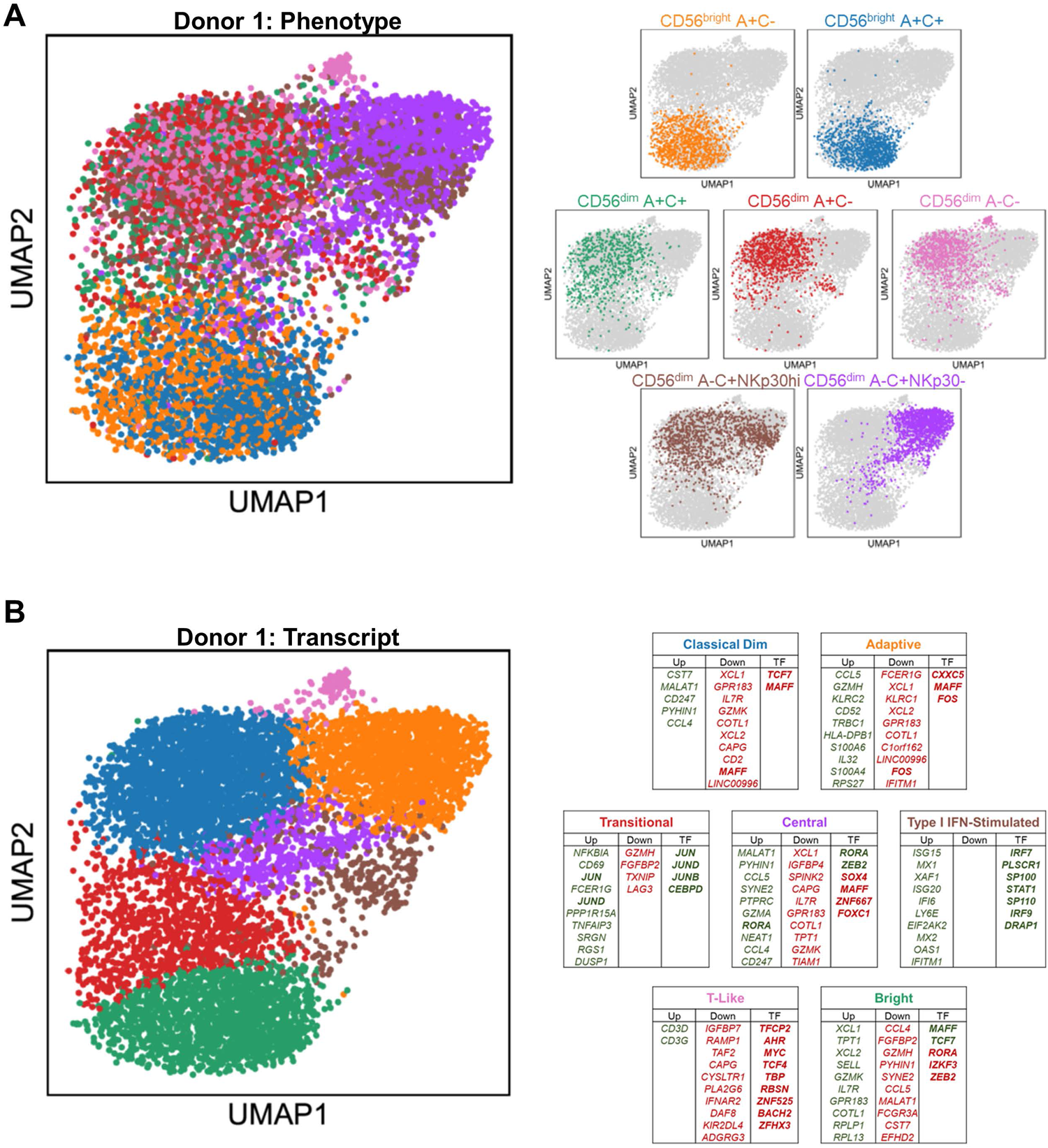
Distinct transcriptional profiles of CD56^bright^, CD56^dim^, and Adaptive NK cells. A) Left, UMAP representation of peripheral blood NK cells from representative donor, color-coded according to phenotypic population hashtag. Right, individual phenotypic populations superimposed on total NK. B) Left, UMAP representation of peripheral blood NK cells from representative donor, color-coded according to transcriptional cluster. Right, top 10 consistently upregulated (green), downregulated (red), and transcription factor (bold) genes among the top 100 DEGS of both donors for each transcriptional cluster; Central and T-Like clusters are unique to this donor.

Five common transcriptional clusters emerge among the peripheral blood NK cells, driven by their composition and their state: Bright, Classical Dim, Adaptive, Transitional Activation, and Type I IFN-stimulated (Figure 2B, S2C). Two additional clusters are present in one donor but not the other: a small T-Like population marked by expression of *CD3D* and *CD3G*, which has been observed in some Adaptive NK^23^; and a centrally located cluster with no obvious distinguishing feature other than higher gene content. Using genes consistently upregulated and downregulated among the top 100 DEGs of the two donors, signatures were defined for each of the transcriptional clusters (Table S1). The Bright cluster, containing much of the CD56^bright^ A+C- and A+C+ populations, express *SELL*, *IL7R*, *GZMK*, the chemokines *XCL1* and *XCL2*, and the transcription factors *MAFF* and *TCF7* (Figure S2D, 2B). The Classical Dim cluster, containing much of the diverse array of CD56^dim^ “non-Adaptive” phenotypic populations, is distinguished by the expression of effector molecules such as *CST7* and *CD247* and the chemokine *CCL4*. The Adaptive cluster, containing all the phenotypic Adaptive and half of the phenotypic “non-Adaptive” populations, distinguishes itself with expression of *GZMH*, *IL32*, *CD52*, the chemokine *CCL5*, and HLA Class II. The Type I IFN-stimulated cluster is composed of all the phenotypic populations in roughly equal contribution, potentially revealing the dramatic effect this early pathogen detection signal has on the transcriptome to overwhelm all other differences. The Transitional Activation cluster, a mix of CD56^bright^ and CD56^dim^ cells, is distinguished by early response factors and negative feedback regulators such as *JUN*, *JUND*, *CEBPD*, and *NFKBIA*. While these cells may be grouped together simply because of their activation status, localization to the Bright and Classical Dim interface and the dearth of Adaptive NK cells suggests a population-specific response and transitional state mediated by AP-1 factors and NFκB signaling. Velocity analysis implicates two possible transcriptional courses, both originating from the Bright cluster: one moves through the Transitional Activation stage onto Classical Dim followed by Adaptive, while the other passes through Type I IFN stimulation and onto Adaptive via a sparsely populated path (Figure S2E).

### Adaptive and CD56^bright^ NK Cells Share a Transcriptional Program Inverse of Classical CD56^dim^ NK Cells

A striking feature of the UMAPs of both donors is the curvature of the structure: the Bright and Adaptive NK cells, although at either end of the contiguous mass of cells, arc toward each other while the Classical Dim NK cells in between them are pushed in the opposite direction. As the Bright cells that were most outstretched toward the Adaptive were the A+C+ population, we considered the possibility that the signal from shared *KLRC2* expression was responsible for this observation. We removed *KLRC2* from the analysis to test this, but the overall structure remains preserved (Figure 3A). In contrast, removal of either the CD56^bright^ or CD56^dim^ A-C+ phenotypic populations collapsed the structure (Figure S3A). This underlying link between the Bright and Adaptive clusters is driven by the transcriptional program within NKG2C+ NK cells, not an artifact of *KLRC2* gene expression itself.

**Figure 3.**
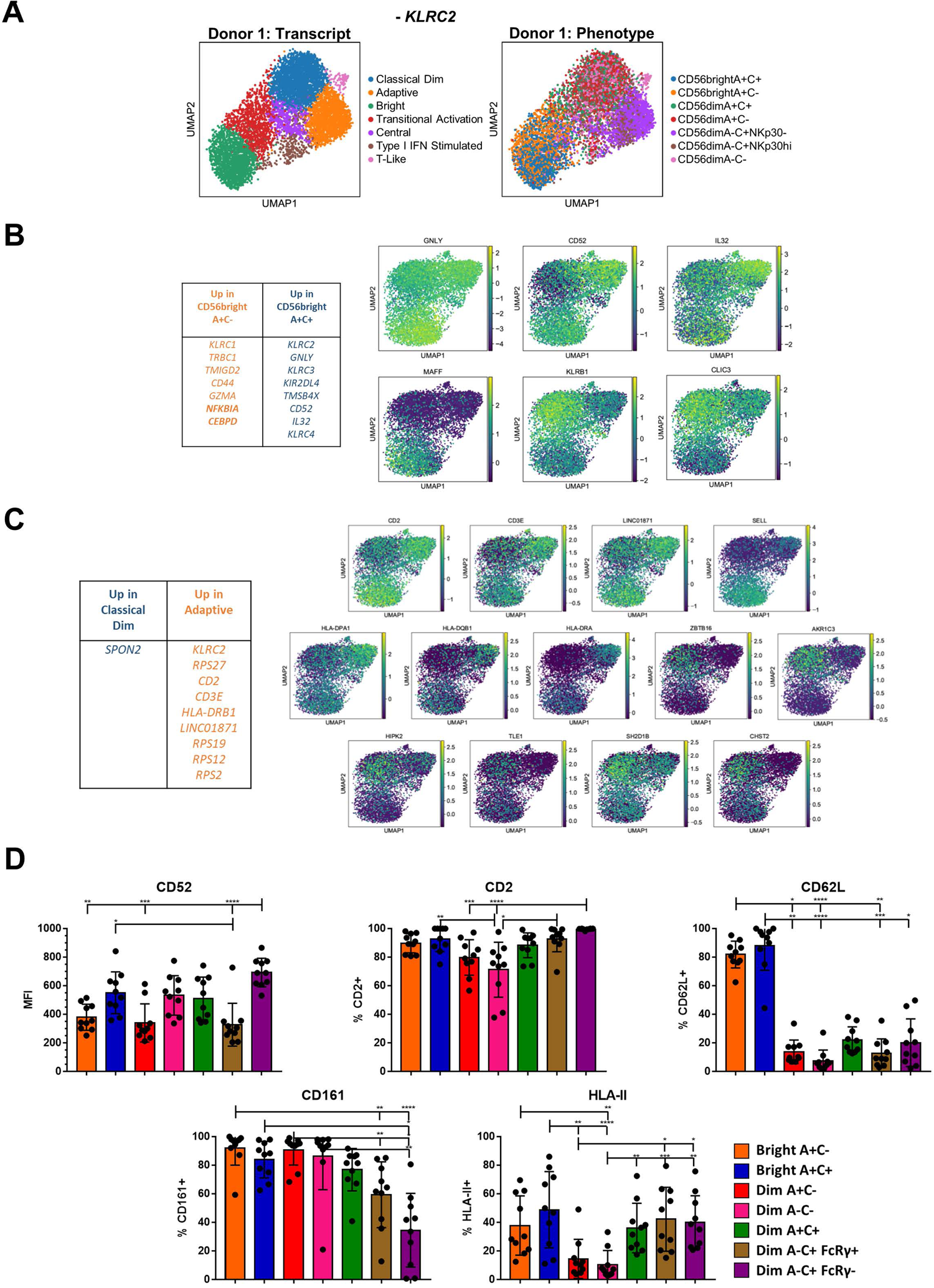
Convergence of a common Bright-Adaptive gene program. A) UMAP of representative donor with *KLRC2* removed from analysis, color-coded according to transcriptional cluster (left) or phenotypic population (right). B) Top DEGs between the CD56^bright^ A+C- and CD56^bright^ A+C+ phenotypic populations. Right, heatmaps of selected DEGs. C) Top DEGs between the Classical Dim and Adaptive transcriptional clusters, heatmaps of selected DEGs above and to the right. D) Surface expression of proteins encoded by genes commonly expressed by Bright and Adaptive transcriptional clusters on NK populations from CMV+ donors. Dunn’s multiple comparison test performed, comparing groups below tick marks to group below capped end.

While CD56^bright^ A+C- and CD56^bright^ A+C+ NK cells cluster together when analyzing the total peripheral blood NK population, a direct comparison of the two phenotypic populations confirmed that the CD56^bright^ A+C+ population was more akin to the Adaptive cells: increased expression of *GNLY*, *CD52*, and *IL32*, and decreased expression of *CLIC3*, *MAFF*, and *KLRB1* (Figure 3B). Similarly, a direct comparison of the Adaptive and Classical Dim transcriptional clusters uncovered additional DEGs shared by the Adaptive and Bright NK cells: increased expression of *CD2*, *CD3E*, *LINC01871*, *SELL*, and HLA Class II, and decreased expression of *HIPK2*, *TLE1*, *SH2D1B*, *ZBTB16*, *CHST2*, and *AKR1C3* (Figure 3C). Several components of this common Bright-Adaptive gene set were validated at the protein level (Figure 3D). Another shared pattern are the ribosomal protein genes: among the over 80 genes, the overwhelming majority were expressed higher in Bright and Adaptive NK cells than in Classical Dim NK cells, despite equivalent gene and transcript count in these clusters (Figure S3B). The canonical phenotypic differences between the Bright and Adaptive NK cells belie the common transcriptional program linking the two, including the preparation for intensive ribosomal biogenesis^25^ – poised to generate an abundance of proteins for effector function, proliferation, or both.

### Adaptive NK Cells are Uniquely Marked by Loss of the Transcription Factor CXXC5, a Marker of Innateness

Memory cells are not simply the transient intermediate between immaturity and terminal differentiation, but a separate and stable state. The single-cell RNAseq data separates the Adaptive NK cell cluster from the other NK cells on a transcriptional basis. While there are many similarities to the Bright cluster, unique to the Adaptive cluster is downregulation of the transcription factor *CXXC5* (Figure 4A). There are no reports of CXXC5 function in NK cells, but in other settings it has been shown to be a co-factor of TET2 and SUV39H1^26–29^ - epigenetic remodelers that repress memory and naive-associated genes in T cells^26,30,31^. Flow staining showed that CXXC5 protein is lower in the NKG2C+ NK population only in CMV+ individuals, and lower levels could be seen in the subset of NKG2A-NK cells with an otherwise Adaptive phenotype (CD161-CD2hi, Figure 3D) in a CMV+ individual with both *KLRC2* genes deleted (Figure 4B-C). Unlike PLZF, which is downregulated in both CD56^bright^ and Adaptive NK cells, decreased CXXC5 is a unique marker of Adaptive NK cells (Figure 4D). In fact, extending the analysis to T cells, we find that CXXC5 protein levels continue to go down from the NK-like NKG2C+ CD56+ T cells to the CD62L-effector populations and finally to the CD62L+ memory and naive T cell populations, which essentially have none (Figure 4E). CXXC5 can therefore be a measure of innateness, with low expression of CXXC5 characteristic of the most adaptive of the NK cells.

**Figure 4.**
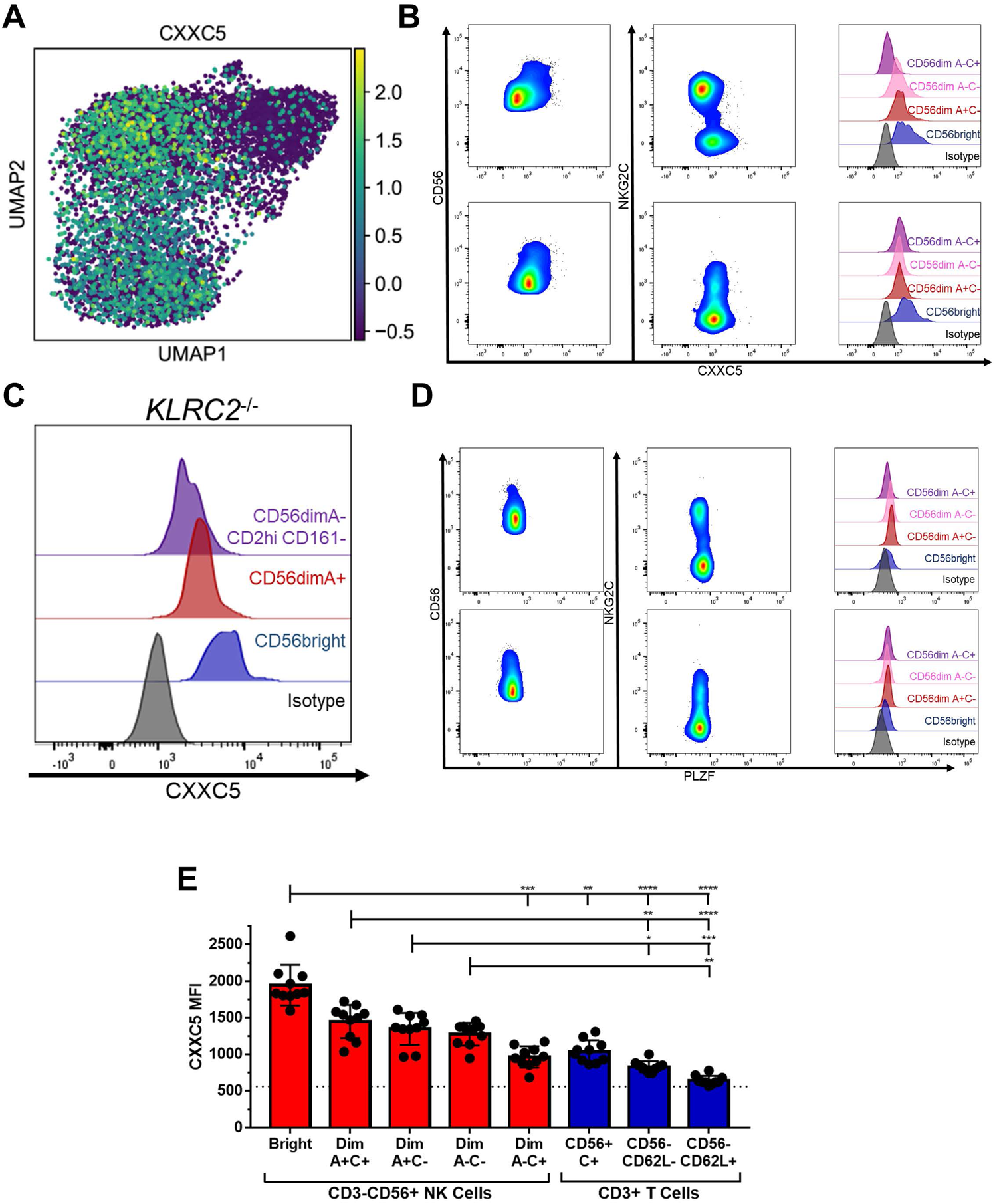
Bright and Adaptive NK Cells are distinguished from each other by expression of CXXC5. A) Heatmap of *CXXC5* expression among Donor 1 NK cells. B) Flow cytometry staining of NK cells for CXXC5 in representative CMV+ (upper) and CMV-(lower) donors. C) Flow cytometry staining of NK cells for CXXC5 in a CMV+ donor with no copies of *KLRC2*. D) Flow cytometry staining of NK cells for PLZF in representative CMV+ (upper) and CMV-(lower) donors. E) Expression of CXXC5 in NK and T cell populations from CMV+ donors; dashed line represents isotype staining. Dunn’s multiple comparison test performed, comparing groups below tick marks to group below capped end.

### Adaptive NK Cells Acquire Superior Effector Function While Retaining the Capacity for Proliferative Burst

The metamorphosis from naive to memory or effector cells is the tradeoff between a cell’s self-renewal capacity and the speed and robustness of its effector response. A fair comparison of NK cell responses to a target is challenging due to the diversity of receptors and ligands expressed heterogeneously across the various NK cell populations^14^. To bypass these superficial biases, we used an agnostic stimulus in the form of treatment with PMA and ionomycin and measured IFNγ production; baseline *IFNG*, unlike the genes for granzymes and cytolytic molecules, was not among the top 100 DEGs for these phenotypic populations or transcriptional clusters. After 5 hours of stimulation, a higher frequency of the CD56^dim^ A-C+ FcRγ-Adaptive population produced IFNγ than any of the other phenotypic populations (Figure 5A). NK cells expressing a self-KIR produced more IFNγ than those not expressing a self-KIR, indicating that the effect of education is not synapse-dependent^24,32^ (Figure 5B). The Adaptive NK cell population typically expresses CD57 and self-KIR but is still superior to the other CD57+KIR+ CD56^dim^ NK cells (Figure 5C). The CD56^bright^ A+C-CD57-KIR-population produced less IFNγ than the equivalent CD56^dim^ A+C-CD57-KIR-population (Figure 5D). The low amount of IFNγ produced by the CD56^bright^ NK cells is surprising given how well they respond to tumor target lines in a co-culture functional assay (Figure S1B). Unlike in the co-culture functional assay, the NK cells in this agnostic stimulus were not pre-treated with IL-2 so as to measure baseline capacity. Performing a time course of PMA and ionomycin challenge revealed that, when pre-treated with IL-2 overnight, the CD56^bright^ NK cells produce IFNγ as early as one hour after stimulus and respond consistently, while the CD56^dim^ NK cells accumulate their IFNγ response over time (Figure 5E). Whereas the CD56^bright^ cells require priming to respond, akin to the necessity of co-stimulation for naive T cells^33,34^, the Adaptive NK cells have no such hindrance.

**Figure 5.**
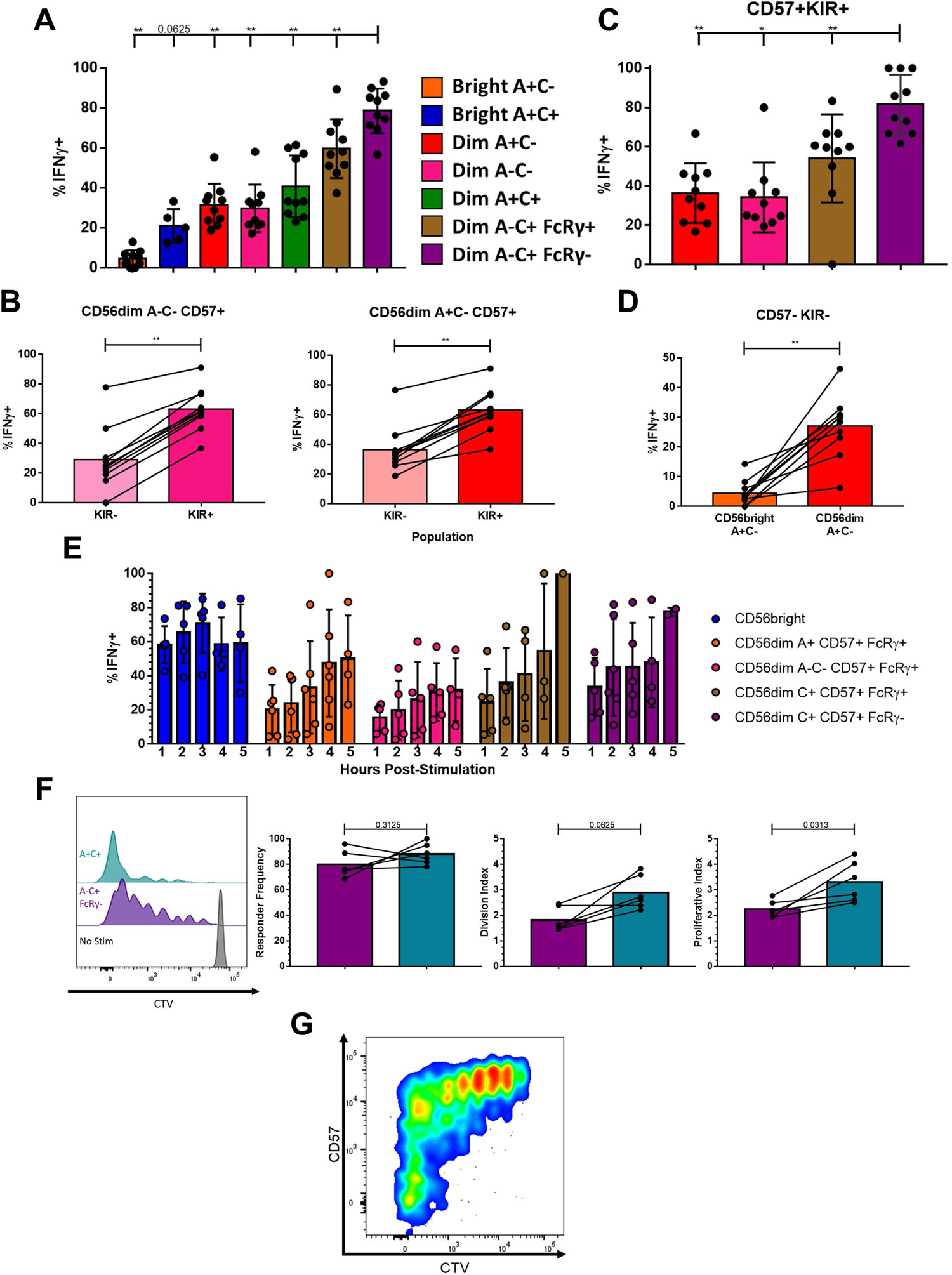
Memory-like effector function and proliferative capacity is intrinsic to Adaptive NK cells. A) IFNγ production in NK cell populations from CMV+ donors following five hour stimulation with PMA+Ionomycin; Wilcoxon test performed on indicated pairs. B) IFNγ production among NK cell populations expressing any one or more of KIR2DL1/2DS1/2DL2/2DS2/2DL3/3DL1/3DS1 or negative for all of these following five hour stimulation; Wilcoxon test performed. C) IFNγ production in NK cell populations expressing CD57 and at least one self-recognizing KIR (KIR2DL2 and/or KIR2DL3) following five hour stimulation; Wilcoxon test performed on indicated pairs. D) IFNγ production among NK cell populations negative for CD57 and KIR2DL1/2DS1/2DL2/2DS2/2DL3/3DL1/3DS1 following five hour stimulation; Wilcoxon test performed. E) IFNγ production in NK cell populations at timepoints indicated following stimulation, after overnight culture with 200U/mL IL-2. F) NK cell populations 9 days after stimulation. Left, dilution of proliferation dye in representative donor shown. Right, proliferation kinetics with lines connecting samples from the same donor; Wilcoxon test performed. G) Expression of CD57 and proliferation dye among A-C+ FcRγ-NK cells nine days after stimulation with PMA+Ionomycin, representative donor shown.

The benefit of Adaptive NK cells and their functional superiority would be lost if they were unable to proliferate and form a long-lived niche. Addressing the same heterogeneity challenge, we stimulated peripheral blood NK cells with a lower dose of PMA and ionomycin. The Adaptive NK cells underwent robust proliferation after nine days of stimulation (Figure 5F); intriguingly, some of the FcRγ-cells that grew the most were CD57-, which are rare among Adaptive cells in circulation (Figure 5G). While Adaptive NK cells initiated cell division at a similar rate to A+C+ NK cells, the A+C+ NK cells underwent an average of one additional round of cell division. Therefore, the Adaptive NK cell population retains proliferative capacity while acquiring superior effector function.

## Discussion

The close transcriptomic and phenotypic linkage between the Bright and Adaptive NK cell populations upends a linear model of human NK cell differentiation, introducing a second axis and branch of development. The CD56^bright^ A+C+ NK cells are central to this due to their increased expression of Adaptive features and Bright-Dim-blurring surface phenotype. The exact tipping point from this population to Adaptive NK cells – the why, where, and how this differentiation step occurs – is a subject critical for further study, both to understand memory formation across innate and adaptive lymphocytes and to generate this functionally superior subset of cells on-demand for therapeutic use.

The co-expression NKG2A and NKG2C on this precursor population is particularly curious – both receptors signal in these cells, but it is unclear if ligation of either or both is central or even relevant to their further differentiation. The presence of otherwise phenotypically Adaptive NK cells in individuals who lack the gene for NKG2C indicates that signaling through this receptor – and recognition of CMV antigens through it – while advantageous, is not absolutely necessary for this. This dispensation of NKG2C would also imply that CMV, while a potent driver of the Adaptive NK population, would not be the sole cause. Nonetheless, the expression of CD62L among both the CD56^bright^ and Adaptive NK offers a tantalizing clue as to where to find this transition in persons undergoing a primary CMV infection: matched single-cell analysis of NK cells in lymphoid organs and peripheral blood would likely provide invaluable data to fill in the missing link.

Overnight culture of NK cells with IL-12, IL-15, and IL-18 has been shown by many groups to markedly improve the function of these “cytokine-induced memory-like” cells^35,36^. These NK cells have a transcriptomic signature resembling CD56^bright^ NK cells^37^, which may explain their conflation with memory/Adaptive NK cells despite the absence of many features of an Adaptive-specific phenotypic signature. Indeed, the many similarities between the Bright and Adaptive NK cells underscore a potential mechanistic role for their differences in delineating naiveness from memory. In one example, both populations express the long intergenic non-coding RNA *LINC01871*, in contrast to its absence in the classical CD56^dim^ population, while the Bright alone express *LINC00996*; the transcriptional program in common between the Bright and Adaptive could be mediated in part by the former, and the differences a consequence of the latter. Loss of the transcription factor CXXC5, a partner of the T cell memory repressors TET2 and SUV39H1^26–29^, in the Adaptive NK populations presents a parsimonious model for the gain of memory features in this population. This is especially the case considering that the Adaptive NK population expresses levels of CXXC5 similar to its TCR-expressing doppelganger, the NKG2C+ CD8+ TCRαβ+ population^38^, placing them shoulder-to-shoulder at the border of innate and adaptive cytotoxic lymphocytes.

The dissociation between the surface phenotype and transcriptome of the NK cell populations is one of the most striking findings of this study. For example, the conglomeration of most CD56^dim^ NK populations as a single transcriptional cluster is in defiance of their phenotypic diversity and even functional capacity. As an example, KIR+ NK cells are intrinsically superior producers of IFNγ than KIR-NK cells, but that advantage may be regulated at the protein level, rather than dependent upon transcriptional shifts. Therefore, the few effector molecule DEGs may point to more fundamental differences in the populations’ roles in the immune response, such as XCL1/2 from CD56^bright^ cells recruiting cross-presenting XCR1+ dendritic cells, or cystatin-F protecting Classical Dim cells from activation of their potent Granzyme B cargo. Especially curious was the co-localization of half the CD56^dim^ A-C+ NKp30hiFcRγ+ NK cells with the Classical Dim and half with the Adaptive NK; the latter had downregulated *FCER1G* expression but still expressed FcRγ protein, a reminder that cell phenotype does not foretell fate.

A major implication of the Bright-Adaptive transition is the possibility of clonal expansion and maintenance among NK cells. In the absence of unique, heritable, and immutable rearranged antigen receptor sequences, clonal expansion has been inferred from strong data showing pools of Adaptive NK cells with shared patterns of chromatin accessibility and mitochondrial DNA somatic mutations within individuals^22^. On the other hand, the inflammatory environment of CMV infection has been shown to stunt the proliferation of Adaptive NK cells in response to IL-2 and IL-15^39^. One hypothesis to reconcile these results is that the signal supporting the outgrowth of this population synergizes with an activating receptor or comes after CMV is under control and the inflammation environment recedes, while another is the possibility of an unexplored “stem memory” population of Adaptive NK cells, such as the CD57-NKG2C+ FcRγ-population that expands in response to stimulus-agnostic PMA and ionomycin. An advantage of preserving an NK “clone” in the absence of a unique peptide-specific receptor is that the diversity of receptor expression across the NK cell compartment creates unique compositions of NK cell functional capacity^14^, including individual cells that may be equipped to be highly effective in certain situations. Thus, a CD56^bright^ “naïve” NK cell expressing the CMV-recognizing receptor NKG2C may be shunted down a parallel track to form a persistent and self-renewing memory population to control a recurrent chronic viral infection.

## Materials & Methods

### Flow Cytometry

Extracellular staining for CD3 (BD, clone UCHT1: BrilliantViolet650), CD56 (Beckman Coulter, clone N901: ECD), CD8 (Biolegend, clone RPA-T8: BrilliantViolet570), NKG2A (Miltenyi, clone REA110: PE-Vio770), NKG2C (Miltenyi, REA205: APC, PE-Vio770, VioBright FITC; R&D, clone 134591: PE), KIR2DL1/S1 (Miltenyi, REA284: FITC), KIR2DL2/L3/S2 (BD, Clone CH-L: BrilliantViolet605, FITC), KIR3DL1/S1 (Miltenyi, REA168: APC-Vio770, FITC), CD57 (BD, clone HNK-1: BrilliantViolet421, PE), CD62L (Biolegend, clone DREG56: BrilliantViolet785; Beckman Coulter, clone DREG56: PE), NKG2D (Biolegend, clone 1D11: BrilliantViolet605), CD16 (BD, clone 3G8: BrilliantViolet711), NKp46 (Biolegend, Clone 9E2: PerCP-Cy5-5), NKp30 (Miltenyi, clone AF29-4D12: PE), CD2 (Biolegend, clone RPA-2.10: BrilliantViolet711), CD52 (Miltenyi, clone REA164: VioBlue), CD161 (Miltenyi, clone REA631: PE), HLA-DR/DP/DQ (Miltenyi, clone REA332: APC-Vio770), CD107a (BD, clone H4A3: BrilliantViolet786), CD14 (Biolegend, clone 63D3: AlexaFluor700; Miltenyi, clone TUK4: FITC), CD4 (Biolegend, clone SK3: AlexaFluor700), TCRVδ1 (Miltenyi, clone REA173 : APC-Vio770), PD-1 (Biolegend, clone EH12.2H7: PacificBlue), KIR3DL1 (BD, clone DX9: BrilliantViolet711), TCRγδ (eBioscience, clone B1.1: PE), KIR2DL1/S1 (Beckman Coulter, clone EB6B: PE-Cy5-5) was performed at room temperature for 30 minutes in PBS (w/o Ca or Mg) with 0.5% BSA and 2mM EDTA. Intracellular staining for IFNγ (BD, clone B27; AlexaFluor700) and FcRγ (Millipore, rabbit polyclonal against FcERI γ subunit; FITC) was performed after fixation and permeabilization with Fix & Perm Cell Permeabilization Reagents (Life Technologies) according to manufacturer’s instructions. Transcription factor staining for CXXC5 (Cell Signaling, clone D1O4P; labeled with anti-rabbit IgG (H+L) F(ab’)^2^ AlexaFluor647, Cell Signaling) and PLZF (eBioscience, clone Mags.21F7; AlexaFluor488) performed using FoxP3/Transcription Factor Staining Buffer Set (eBioscience) according to manufacturer’s instructions. Viability was assessed by staining with DAPI (Sigma) or Live-Dead Fixable Aqua (Invitrogen). Analysis of flow data was performed in FlowJo 10.8.0 (BD).

### Cell Culture

PBMCs were isolated from buffy coats obtained from healthy volunteer donors via the New York Blood Center (NYBC). The MSKCC Institutional Review Board (IRB) waived the need for additional research consent for anonymous NYBC samples. Human donors *KLRC2* copy number was determined by PCR^40^. PBMC were cryopreserved in fetal bovine serum with 10% DMSO. K562, 721.221, and BE(2)n (provided by Dr. Nai-Kong Cheung) cells were tested regularly for mycoplasma via PCR. The HLA-E:G*01 construct was designed and expressed in K562s as described previously^38^; the HLA-E:A*02 construct (leader sequence MAVMAPRTLVLLLSGALALTQTWA) from Genscript was designed and expressed in a similar fashion^38^. All cells were cultured in RPMI 1640 with 10% FCS and Penicillin-Streptomycin.

### Patients and Transplant Procedures

Patients and donors provided informed written consent for research, and studies were approved by the MSKCC Institutional Review Board. Of 267 total adult patients receiving HCT at MSKCC between 2006 to 2017 as part of a larger study^19^, 20 patients who received a CD34+ selected graft (CliniMACS, Miltenyi) were analyzed further and included here. Patients did not receive additional pharmacologic GVHD prophylaxis. All patients received acyclovir prophylaxis starting on admission for HCT and continued for at least 12 months, and recipients who were CMV+ or CMV-with a CMV+ donor were routinely monitored at least weekly starting on D14 post-HCT. CMV infection/reactivation was determined by pp65 antigenemia assays prior to 2010 and by CMV qPCR (Roche Diagnostics) from 2010 onward.

### Functional Assays

Degranulation of NK cells was assessed by labeling with CD107a antibody in culture. IFNγ production was measured by blocking Golgi transport with brefeldin A (MP Biomedicals) and GolgiStop (BD) 1 hour after stimulation, followed by intracellular staining four hours later unless otherwise indicated. Replicates with fewer than 100 cells in the selected populations excluded.

Co-culture with tumor cell targets was performed at a ratio of 1:1 PBMCs to Target cells in 100μL. ADCC assays were performed with BE(2)n cells labeled with 1μg/mL of the anti-GD2 clone 3F8 (provided by Dr. Nai-Kong Cheung). IFNγ production was also assessed following stimulation with 50ng/mL PMA (Sigma) and 1μg/mL ionomycin (MP Biomedicals). PBMCs were pre-treated with 200 U/mL of IL-2 (Peprotech) overnight as indicated.

For proliferation assays, PBMC were labeled with (1.66μM) CTV and plated 250,000 per well and rested overnight, before stimulation with 0.625ng/mL PMA and 0.5μg/mL ionomycin.

### Single-Cell RNA Sequencing

PBMCs from two CMV+ donors with two intact alleles of *KLRC2* were thawed and sorted as shown in Fig S2A. Sorted populations were labeled with TotalSeq-B Anti-Human Hashtag antibodies (Biolegend), and 3,000 cells from each of the seven sorted populations were combined for each donor and submitted to MSKCC’s Integrated Genomic Operation core facility for 10x Genomics library prep and 3’ Gene Expression and 5’ Feature Barcode sequencing. The software Cell Ranger (version 5.0.1; 10x Genomics) was used for demultiplexing and alignment using default parameters. Velocyto (version 0.17.17) performed counting of spliced and un-spliced RNA molecules. Pre-processing and quality control of the data was carried out using the Python software package SCANPY (version 1.8.1). Cells that expressed low gene numbers, reading depth (counts) or high mitochondrial or ribosomal gene percentage were removed, the borders for these parameters were adapted according to the experiment. Cell cycle and mitochondrial genes were regressed out. SCANPY was also used to perform dimensionality reduction and clustering. The neighborhood graphs were based on 40 principal components and 80 neighbors. Clustering was performed using the Leiden algorithm with resolution r = 1.1. UMAP dimensionality reduction was computed using SCANPY’s default parameters. The software tool scvelo (version 0.2.2) was used to compute RNA velocities within a deterministic model.

### Statistical Analysis

Statistical analyses were performed in GraphPad Prism 7.00 as described in the legends. Bars represent mean and SD; * p <0.05, ** p <0.01, *** p <0.001, **** p<0.0001.

## Data Availability

Raw and processed sequencing files are deposited with GEO under the Accession Number GSE243030.

## Author Contributions (per CRediT Taxonomy)

Conceptualization: MKP and KCH.

Formal Analysis: MKP and SG.

Funding: KCH.

Investigation: MKP, RS, J-BLL, TK, and KvdP.

Methodology: MKP and SG.

Supervision: KCH and JCS.

Writing – Original Draft: MKP.

Writing – Reviewing and Editing: All authors.

## Acknowledgements

MKP and KCH are inventors on a patent application for the design and use of HLA-E:peptide chimeric molecules. The authors would like to thank Drs. Kyle B. Lupo, Gianluca Scarno, and Sanam Shahid for thoughtful feedback on the manuscript.

These studies were funded by NIH R01 AI150999 and U01 AI069197, and the MSKCC Human Oncology and Pathogenesis Program. SG is the recipient of a CRI/Donald J. Gogel Fellowship (CRI #3934).

---

**Supplemental Figure 1.**
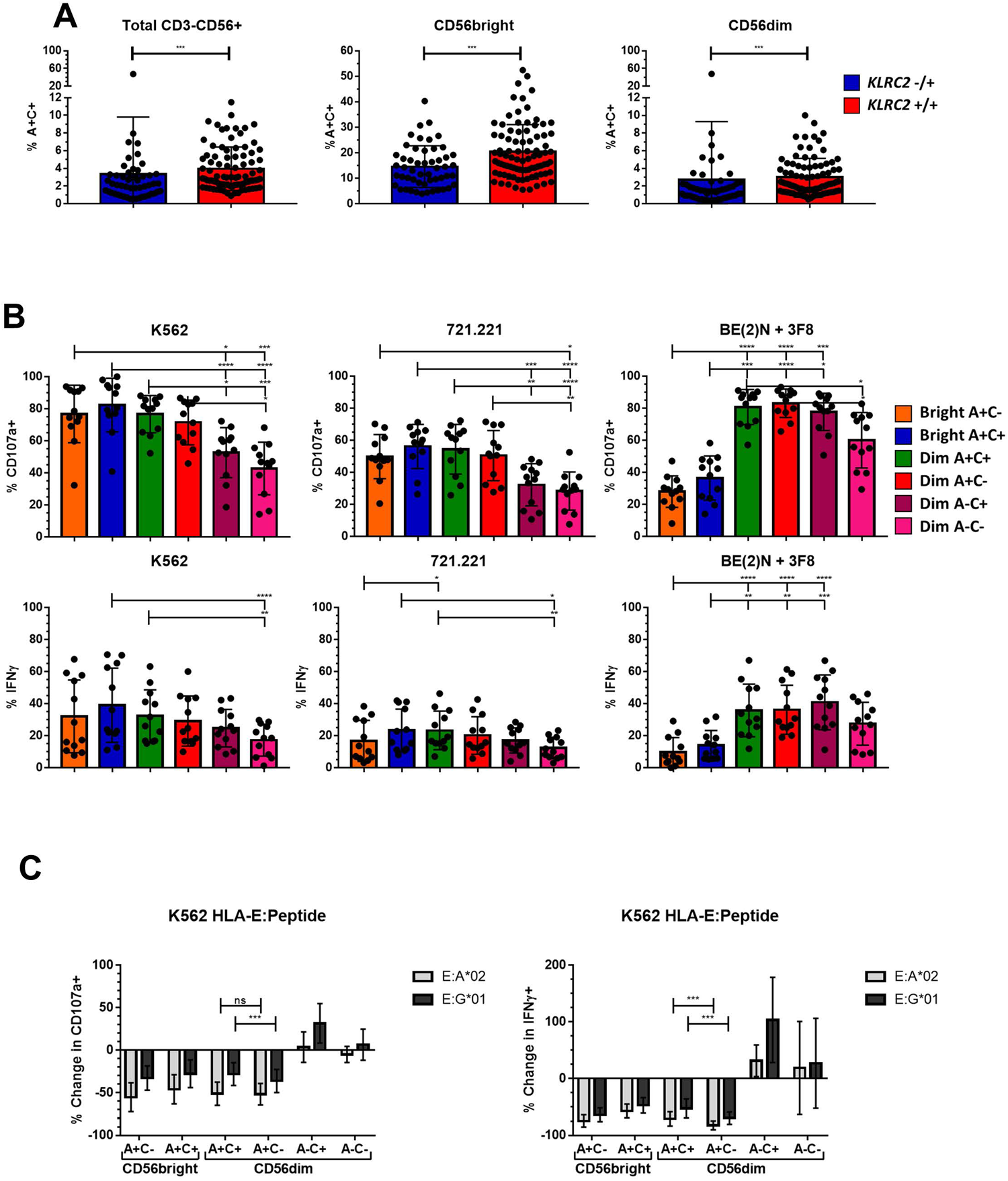
Genetic and functional characteristics of the immature NKG2C+ precursor for adaptive NK cells. A) Top, frequency of A+C+ among indicated populations according to *KLRC2* copy number; Mann-Whitney test performed. B) Degranulation and IFNγ production of NK cell populations after 5 hour co-culture with the indicated tumor cell lines. Dunn’s multiple comparison test performed, comparing groups below tick marks to group below capped end. C) Percentage change in degranulation and IFNγ production of NK cell populations in co-culture with K562s expressing HLA-E:peptide complexes relative to non-transduced K562. Wilcoxon test performed on indicated pairs.

**Supplemental Figure 2.**
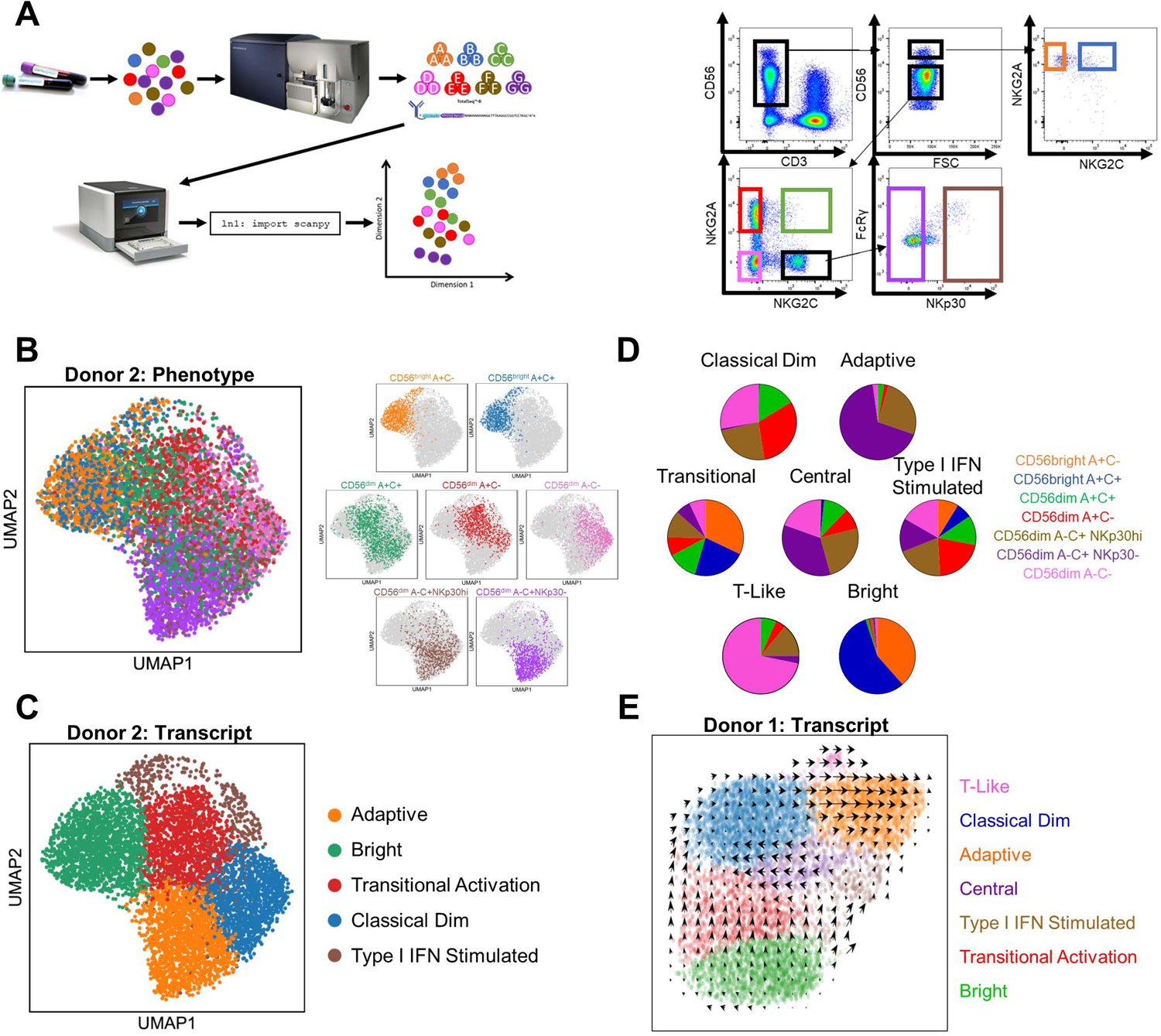
Single-cell transcriptional profiling of phenotypically distinct NK cell populations. A) Left, schematic of single-cell RNA sequencing experimental design. Right, sorting strategy for NK cell populations as represented on one of the two healthy donors. B) Left, UMAP representation of peripheral blood NK cells from second donor, color-coded according to phenotypic population hashtag. Right, individual phenotypic populations superimposed on total NK. C) Left, UMAP representation of peripheral blood NK cells from second donor, color-coded according to transcriptional cluster. D) Phenotypic composition of transcriptional clusters from Donor 1. E) RNA velocity analysis of Donor 1, color-coded according to transcriptional cluster.

**Supplemental Figure 3.**
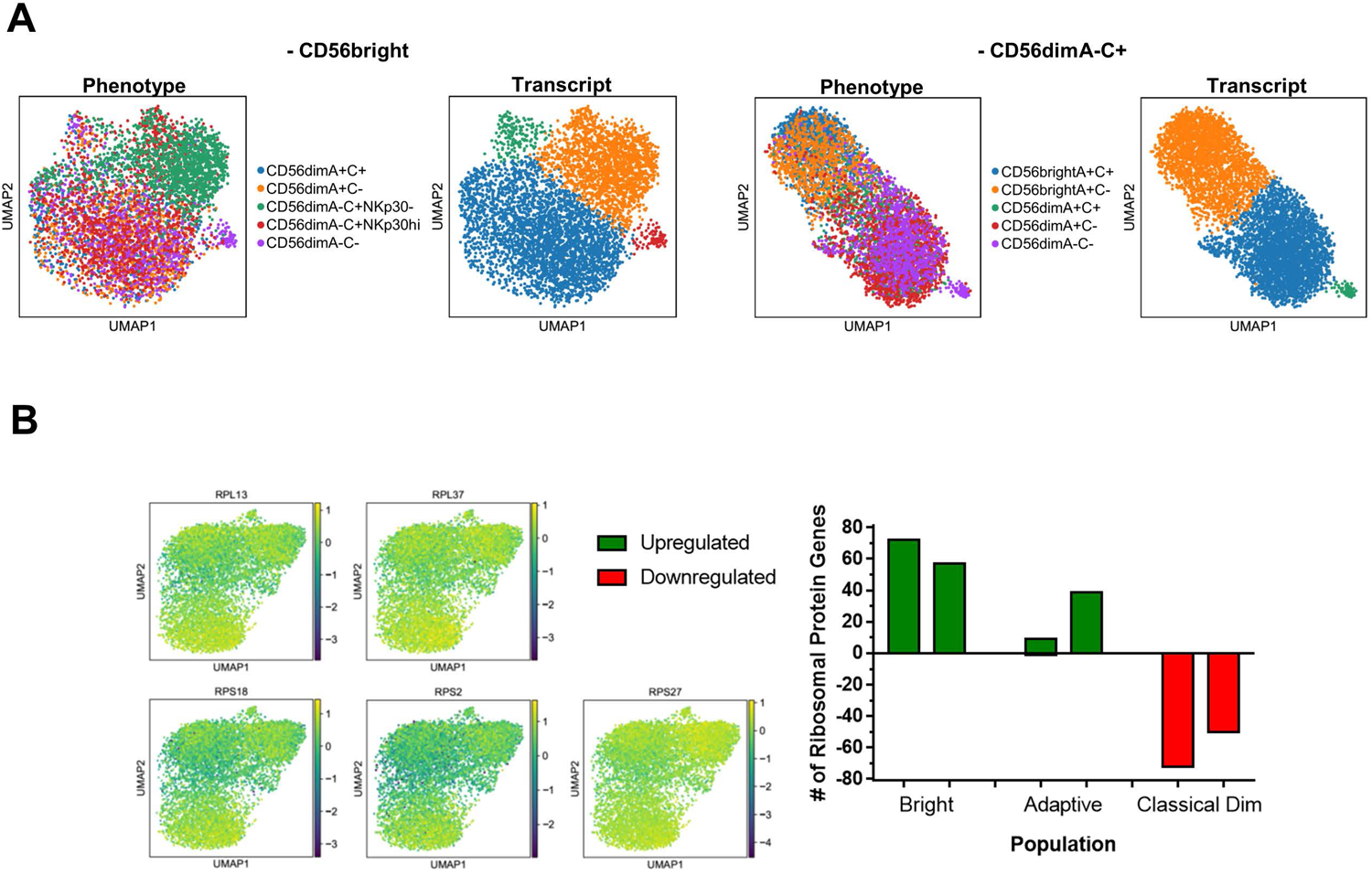
Trancriptomic profile shared by NKG2C+ CD56^bright^ and Adaptive NK cells. A) UMAP of representative donor after removal of phenotypic CD56^bright^ cells (left) or A-C+ cells (right), color-coded according to phenotype (left) or transcriptional cluster (right). B) Left, heatmaps of selected ribosomal protein DEGs. Right, number of genes for ribosomal proteins upregulated or downregulated in the populations of individual donors.

**Table S1.** DEGs by transcriptional cluster and by comparison.

